# Regulation of amino acid and nucleotide metabolism by crustacean hyperglycemic hormone in the muscle and hepatopancreas of the crayfish *Procambarus clarkii*

**DOI:** 10.1101/736603

**Authors:** Wenfeng Li, Kuo-Hsun Chiu, Chi-Ying Lee

**Author notes:** to whom correspondence should be addressed (C-Y L) Chi-Ying Lee.

## Abstract

To comprehensively characterize the metabolic roles of crustacean hyperglycemic hormone (CHH), metabolites in two CHH target tissues of the crayfish *Procambarus clarkii*, whose levels were significantly different between CHH-silenced and saline-treated control animals, were analyzed using bioinformatics tools provided by an on-line analysis suite (MetaboAnalyst). Analysis with Metabolic Pathway Analysis (MetPA) indicated that in the muscle Glyoxylate and dicarboxylate metabolism, Nicotinate and nicotinamide metabolism, Alanine, aspartate and glutamate metabolism, Pyruvate metabolism, and Nitrogen metabolism were significantly affected by silencing of CHH gene expression at 24 hours post injection (hpi), while only Nicotinate and nicotinamide metabolism remained significantly affected at 48 hpi. In the hepatopancreas, silencing of CHH gene expression significantly impacted, at 24 hpi, Pyruvate metabolism and Glycolysis or gluconeogenesis, and at 48 hpi, Glycine, serine and threonine metabolism. Moreover, analysis using Metabolite Set Enrichment Analysis (MSEA) showed that many metabolite sets were significantly affected in the muscle at 24hpi, including Ammonia recycling, Nicotinate and nicotinamide metabolism, Pyruvate metabolism, Purine metabolism, Warburg effect, Citric acid cycle, and metabolism of several amino acids, and at 48 hpi only Nicotinate and nicotinamide metabolism, Glycine and serine metabolism, and Ammonia recycling remained significantly affected. In the hepatopancreas, MSEA analysis showed that Fatty acid biosynthesis was significantly impacted at 24 hpi. Finally, in the muscle, levels of several amino acids decreased significantly, while those of 5 other amino acids or related compounds significantly increased in response to CHH gene silencing. Levels of metabolites related to nucleotide metabolism significantly decreased across the board at both time points. In the hepatopancreas, the effects were comparatively minor with only levels of thymine and urea being significantly decreased at 24 hpi. The combined results showed that the metabolic effects of silencing CHH gene expression were far more diverse than suggested by previous studies that emphasized on carbohydrate and energy metabolism. Based on the results, metabolic roles of CHH on the muscle and hepatopancreas were summarized and discussed.

## Introduction

Crustacean hyperglycemic hormone (CHH) is a polypeptide hormone originally identified in the X-organ/sinus gland (XO/SG) complex of the eyestalks [1-2]. Early biochemical studies of CHHs isolated from various decapod crustaceans showed they are of 72–73 amino acid residues in length, with 3 disulfide bridges formed by 6 highly position-conserved cysteine residues [3-9]; peptide sequencing analyses indicated that CHH is N- and C-terminally blocked; the C-terminal amide was found critical for its hyperglycemic activity [10-14]. These and subsequent molecular studies [15-17] have established that CHH, being a prototypical member, belong to a family of polypeptide hormones, the CHH family, which also includes molt-inhibiting hormone (MIH), vitellogenesis-inhibiting hormone (VIH), and mandibular organ-inhibiting hormone (MOIH), and insect ion transport peptide (ITP) [18-21].

Combined data indicated that CHH is functionally involved with regulation of carbohydrate metabolism and responsible for mediating stress-induced hyperglycemia [22-27]. Specially, glycogen mobilization in two CHH target tissues (the hepatopancreas and muscle) has been shown to be stimulated by CHH (or tissue extracts containing CHH) through regulating activity of the enzymes involved in glycogen metabolism [28-32]; other studies suggested that CHH also stimulates glycolytic flux by increasing the availability of glycolysis substrate (glucose) [22, 33].

In the previous study of ours [34], a double-stranded RNA (dsRNA) technique was employed to effectively and specifically knock down CHH gene expression in the crayfish *Procambarus clarkii*. Metabolites in the tissue (the hepatopancreas and muscle) obtained from the CHH dsRNA-injected (DSI) and saline injected (SAI) animals were detected, identified, and quantified using a nuclear magnetic resonance (NMR)-based metabolomic method. Initial analysis of metabolites directly related to carbohydrate and energy metabolism suggested that CHH regulates tissue metabolism in tissue-specific and complementary manners [34]. In the present study, to fully characterize the effects of silencing CHH gene expression and understand the metabolic roles of CHH, we utilized web-based bioinformatics tools provided by MetaboAnalyst to comprehensively analyze all tissue metabolites whose levels changed significantly after CHH gene silencing. Results not only confirmed our previous conclusion that CHH regulates carbohydrate and energy metabolism but also extended the scope of CHH regulatory roles to amino acid and nucleotides metabolism. Finally, a complete profile of CHH metabolic function is presented and discussed.

## 1. Materials and Methods

### 2.1 Experimental animals and procedures

Extracts of tissues harvested from experimental animals *P. clarkii* untreated, or treated with either saline injection (SAI) or CHH dsRNA injection (CHH DSI) were subjected to ^1^H Nuclear magnetic resonance (NMR) analysis. Experimental treatment protocol, efficacy and specificity of gene silencing by CHH dsRNA, as well as NMR setting parameters, data acquisition and processing, and statistical analysis (Student’s t-test) have been described in a previous study [34].

### 2.2 Metabolic Pathway Analysis (MetPA)

Tissue metabolites whose levels were significantly different between the SAI and CHH DSI groups were subsequently analyzed using resources provided by a web-based tool suite MetaboAnalyst (v. 4.0, http://www.metaboanalyst.ca), specifically two functional analysis modules - Metabolic Pathway Analysis (MetPA) and Metabolite Set Enrichment Analysis (MSEA) [35-37]. Both approaches work by comparing the significant compounds identified from the data uploaded by users to pre-defined functional groups [35].

MetPA combines statistical enrichment analysis with pathway topological characteristics to identify the most relevant pathways under the study condition [36]. Briefly, the metabolites whose tissue levels were significantly different between SAI and CHH DSI groups were uploaded. After data processing and compound name mapping, pathway analysis was performed using the fruit fly (*Drosophila melanogaster*) library from the KEGG database [38]. The hypergeometric test was chosen to be pathway analysis algorithm for the over representation analysis and the relative betweenness centrality for pathway topology analysis.

### 2.3 Metabolite Set Enrichment Analysis (MSEA)

Tissue metabolites whose levels were significantly different between the SAI and CHH DSI groups were subsequently analyzed using resources provided by a web-based tool suite MetaboAnalyst (v. 4.0, http://www.metaboanalyst.ca), specifically two functional analysis modules - Metabolic Pathway Analysis (MetPA) and Metabolite Set Enrichment Analysis (MSEA) [39, 40]. Both approaches work by comparing the significant compounds identified from the data uploaded by users to pre-defined functional groups [35].

MetPA combines statistical enrichment analysis with pathway topological characteristics to identify the most relevant pathways under the study condition [36]. Briefly, the metabolites whose tissue levels were significantly different between SAI and CHH DSI groups were uploaded. After data processing and compound name mapping, pathway analysis was performed using the fruit fly (*Drosophila melanogaster*) library from the KEGG database [38]. The hypergeometric test was chosen to be pathway analysis algorithm for the over representation analysis and the relative betweenness centrality for pathway topology analysis.

The Metabolite Set Enrichment Analysis (MSEA) is a tool to test if there are some biologically meaningful groups (e.g., pathways) of metabolites that are significantly enriched. After data processing and compound name mapping, enrichment analysis was performed using the Overrepresentation Analysis (ORA) against a knowledge database, the Pathway-associated metabolite sets library, which contains 99 metabolite sets based on normal metabolic pathways, in the Small Molecular Pathway Database (SMPDB) [39].

## 2. Results

Our previous study reported that levels of 149 and 181 metabolites in the muscle at 24 and 48 hpi (hour post injection), respectively, and those of 24 and 12 metabolites in the hepatopancreas at 24 and 48 hpi, respectively, were found significantly different [34], between the saline-injected (SAI) and CHH double-stranded RNA-injected (CHH DSI) animals (*Procambarus clarkii*). Among those significantly changed metabolites, those that are directly related to carbohydrate and energy metabolism have been analyzed [34]. In this further study, to give a more comprehensive survey of the effects of silencing CHH gene expression on tissue metabolism, all significantly changed metabolites were analyzed using MetPA (Metabolic Pathway Analysis) and MSEA (Metabolite Set Enrichment Analysis) with web-based resource (MetaboAnalyst, v. 4.0).

### 3.1 MetPA

Analysis with MetPA indicated that the following pathways were significantly affected by silencing of CHH gene expression at 24 hpi in the muscle (Table 1; Supplementary Table1): Glyoxylate and dicarboxylate metabolism, Nicotinate and nicotinamide metabolism, Alanine, aspartate and glutamate metabolism, Pyruvate metabolism, and Nitrogen metabolism. At 48 hpi, only Nicotinate and nicotinamide metabolism remained significantly affected (Table 1; Supplementary Table1).

**Table 1.** KEGG metabolic pathways significantly impacted in the muscle of the crayfish *Procambarus clarkii* by silencing of CHH gene expression as revealed by Metabolic Pathway Analysis.

| Time point | Pathway | Total<sup>a</sup> | Hits<sup>b</sup> | p-Value<sup>c</sup> |
| --- | --- | --- | --- | --- |
| 24 hpi | Glyoxylate and dicarboxylate metabolism | 16 | 7 | 1.9E-03 |
|  | Nicotinate and nicotinamide metabolism | 9 | 5 | 2.5E-03 |
|  | Alanine, aspartate and glutamate metabolism | 23 | 7 | 1.9E-02 |
|  | Pyruvate metabolism | 24 | 7 | 2.4E-02 |
|  | Nitrogen metabolism | 7 | 3 | 0.047 |
| 48 hpi | Nicotinate and nicotinamide metabolism | 9 | 5 | 0.005 |

<sup>a</sup>Total genes in the biological pathway.
<sup>b</sup>Number of genes hit in the biological pathway.
<sup>c</sup>p-Value calculated from raw data.

In the hepatopancreas, silencing of CHH gene expression significantly impacted Pyruvate metabolism and Glycolysis or gluconeogenesis at 24 hpi, and Glycine, serine and threonine metabolism at 48 hpi (Table 2; Supplementary Table2).

**Table 2.** KEGG metabolic pathways significantly impacted in the hepatopancreas of the crayfish *Procambarus clarkii* by silencing of CHH gene expression as revealed by Metabolic Pathway Analysis.

| Time point | Pathway | Total<sup>a</sup> | Hits<sup>b</sup> | p-Value<sup>c</sup> |
| --- | --- | --- | --- | --- |
| 24 hpi | Pyruvate metabolism | 24 | 3 | 0.010 |
|  | Glycolysis or Gluconeogenesis | 25 | 3 | 0.011 |
| 48 hpi | Glycine, serine and threonine metabolism | 25 | 2 | 0.019 |

<sup>a</sup>Total genes in the biological pathway.
<sup>b</sup>Number of genes hit in the biological pathway.
<sup>c</sup>p-V alue calculated from raw data.

### 3.2 MSEA

Analysis using MSEA showed that several Small Molecular Pathway Database (SMPDB) metabolite sets were significantly affected in the muscle at 24 hpi (Table 3; Supplementary Table3), including Ammonia recycling, Nicotinate and nicotinamide metabolism, Aspartate metabolism, Pyruvate metabolism, Alanine metabolism, Glutamate metabolism, Purine metabolism, Arginine and Proline metabolism, Glycine and serine metabolism, Warburg effect, and Citric acid cycle (Table 3; Supplementary Table3). At 48 hpi, the metabolic groups that remained significantly affected included Nicotinate and nicotinamide metabolism, Glycine and serine metabolism, and Ammonia recycling (Table 3; Supplementary Table3).

**Table 3.** SMPDB metabolite sets significantly impacted in the muscle of the crayfish *Procambarus clarkii* by silencing of CHH gene expression as revealed by Metabolite Set Enrichment Analysis.

| Time point | Metabolite set | Total<sup>a</sup> | Hits<sup>b</sup> | p-Value<sup>c</sup> |
| --- | --- | --- | --- | --- |
| 24 hpi | Ammonia Recycling | 32 | 13 | 1.96E-04 |
|  | Nicotinate and Nicotinamide Metabolism | 37 | 14 | 2.7E-04 |
|  | Aspartate Metabolism | 35 | 12 | 0.002 |
|  | Pyruvate Metabolism | 48 | 14 | 0.005 |
|  | Alanine Metabolism | 17 | 7 | 0.006 |
|  | Glutamate Metabolism | 49 | 14 | 0.006 |
|  | Purine Metabolism | 74 | 18 | 0.012 |
|  | Arginine and Proline Metabolism | 53 | 14 | 0.013 |
|  | Glycine and Serine Metabolism | 59 | 15 | 0.015 |
|  | Warburg Effect | 58 | 14 | 0.029 |
|  | Citric Acid Cycle | 32 | 9 | 0.030 |
| 48 hpi | Nicotinate and Nicotinamide Metabolism | 37 | 14 | 0.002 |
|  | Glycine and Serine Metabolism | 59 | 19 | 0.003 |
|  | Ammonia Recycling | 32 | 12 | 0.005 |

<sup>a</sup>Total genes in the biological pathway.
<sup>b</sup>Number of genes hit in the biological pathway.
<sup>c</sup>p-Value calculated from raw data.

In the hepatopancreas, MSEA analysis showed that Fatty acid biosynthesis was significantly impacted at 24 hpi (Table 4; Supplementary Table4).

**Table 4.** SMPDB metabolite sets significantly impacted in the hepatopancreas of the crayfish *Procambarus clarkii* by silencing of CHH gene expression as revealed by Metabolite Set Enrichment Analysis.

| Time point | Metabolite set | Total<sup>a</sup> | Hits<sup>b</sup> | p-Value<sup>c</sup> |
| --- | --- | --- | --- | --- |
| 24 hpi | Fatty Acid Biosynthesis | 35 | 3 | 0.045 |

<sup>a</sup>Total genes in the biological pathway.
<sup>b</sup>Number of genes hit in the biological pathway.
<sup>c</sup>p-Value calculated from raw data.

### 3.3 Changes in amino acid and nucleotide metabolic profiles

Combined results derived from analyses of MetPA and MSEA indicated that silencing of CHH gene expression significantly affected metabolism of carbohydrates (in the muscle and hepatopancreas), nucleotides and amino acids (mainly in the muscle), and fatty acids (in the hepatopancreas). A comprehensive discussion on carbohydrate and energy metabolism in the muscle and hepatopancreas has been reported previously [34]. Here, changes in the profile of metabolites in amino acid and nucleotide metabolism will be described in details.

In the muscle at 24 hpi, levels of several amino acids, including alanine, arginine, aspartate, glutamate, glutamine, proline and serine decreased significantly, while those of 5 amino acids or related compounds, asparagine, anserine, histidine, histamine, and β-alanine, significantly increased (Fig 1). At 48hpi, the changes were similar to those at 24 hpi, with aspartate no longer significantly decreased and anserine no longer significantly increased (Fig 1).

**Figure 1.**
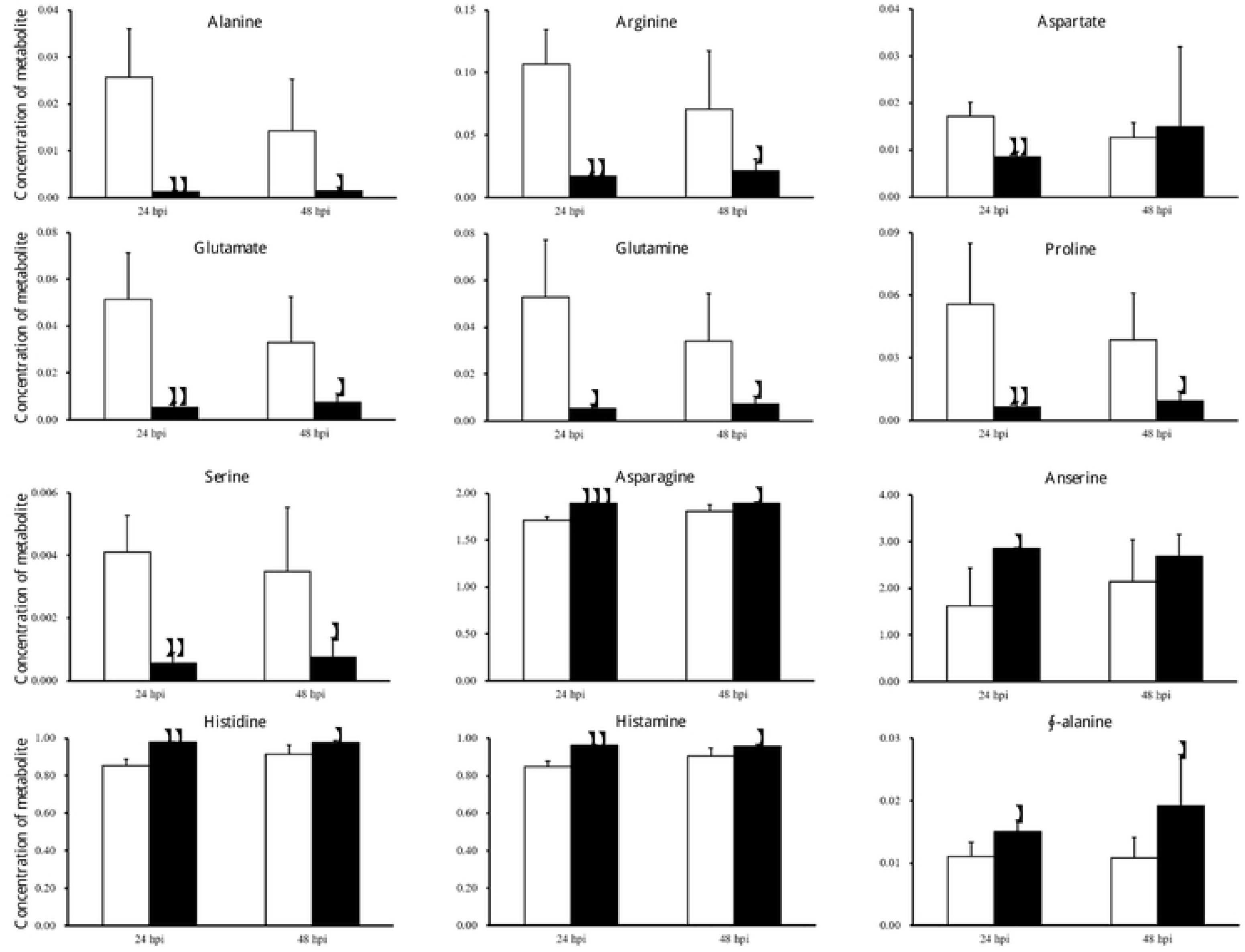
Amino acids and related compounds whose concentration significantly changed by silencing of CHH gene expression in the muscle of the crayfish *Procambarus clarkii* at 24 and 48 hpi. White bar, saline-injected group (SAI); black bar, CHH double-stranded RNA-injected group (CHH DSI).*, p<0.05; **, p<0.01; ***, p<0.001.

In response to silencing of CHH gene expression in the muscle at 24 hpi, metabolite levels involved in nucleotide metabolism significantly decreased almost across the board, including adenine, adenosine, ADP, AMP, ATP, GTP, guanosine, hypoxanthine, inosine, xanthine, xanthosine, cytidine, cytosine, dCTP, dTTP, thymidine, uracil, urea and uridine (Fig. 2). Similar trends of changes were observed at 48 hpi (Fig. 2).

**Figure 2.**
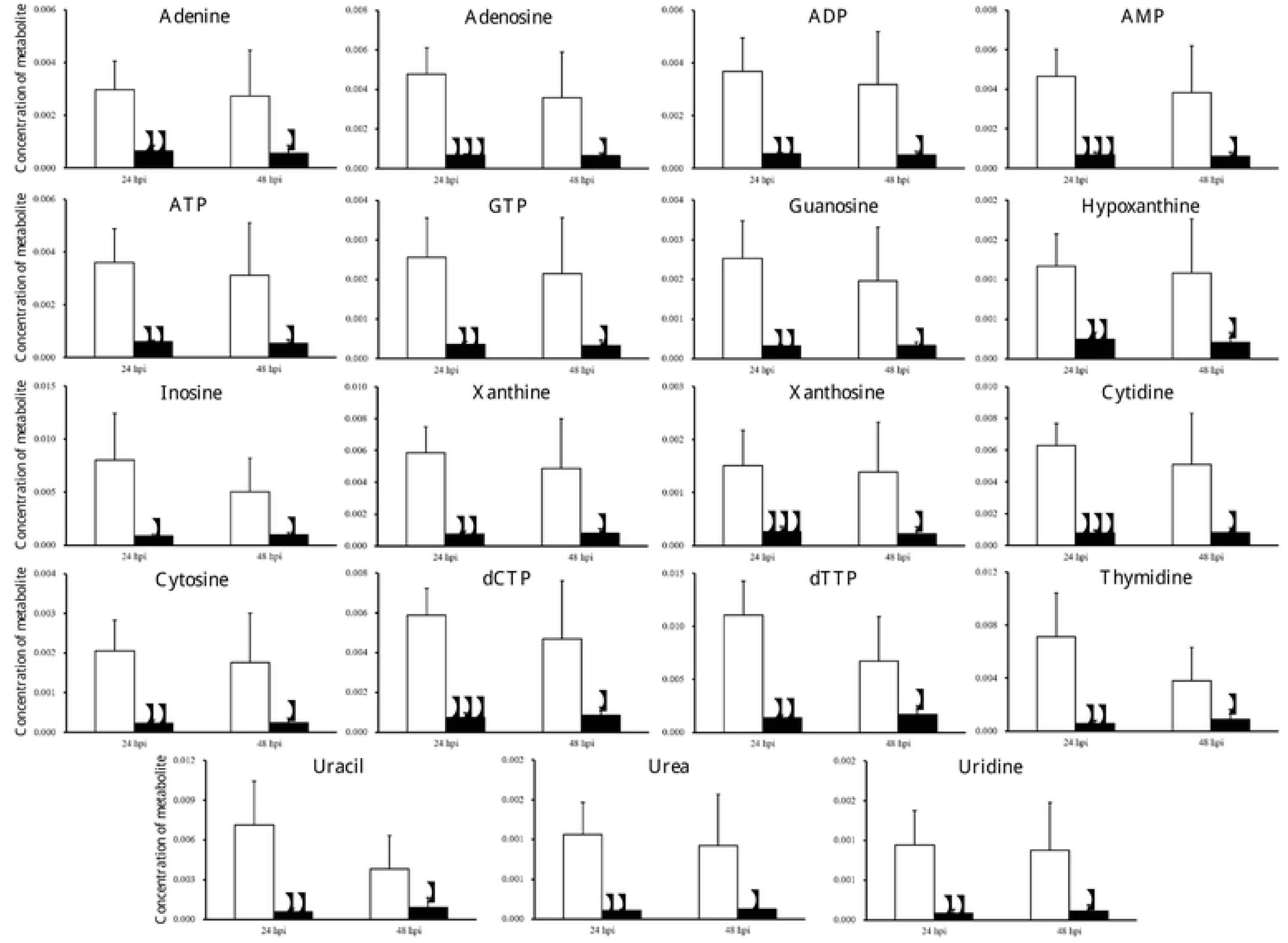
Metabolites involved in nucleotide metabolism whose concentration significantly changed by silencing of CHH gene expression in the muscle of the crayfish *Procambarus clarkii*. White bar, saline-injected group (SAI); black bar, CHH double-stranded RNA-injected group (CHH DSI).*, p<0.05; **, p<0.01; ***, p<0.001.

In the hepatopancreas, with regard to nucleotide metabolism only levels of thymine and urea were significantly decreased at 24 hpi and no significant change was found at 48 hpi.

## 3. Discussion

In the previous study of ours, the significantly changed metabolites related to carbohydrate and energy metabolism have been discussed: under silenced CHH gene expression, the muscle was metabolically characterized as having lower rate of carbohydrates utilization through glycolysis and TCA cycle, and greatly diminished levels of ATP due primarily to reduced synthesis of NAD^+^ and adenine ribonucleotides, whereas the hepatopancreas by unaffected levels of ATP, lower rate of carbohydrates utilization through glycolysis, and increased levels of ketone bodies presumably through enhanced β-oxidation of fatty acids [34].

In this study, comprehensive analysis of the metabolite whose levels were significantly changed by silencing CHH gene expression revealed that the effects were not limited to carbohydrate and energy metabolism as previous studies had indicated [22, 33, 34]. Analysis of the metabolite whose levels were significantly changed by silencing CHH gene expression revealed that the effects were not limited to carbohydrate and energy metabolism as previous studies had indicated. Thus, combined data derived from MetPA and MSEA revealed additionally that, in the muscle nucleotide metabolism was severely reduced, likely due to a negatively impacted pentose phosphate pathway (PPP), and amino acid metabolism significantly affected that would likely result in slowing down protein synthesis, while in the hepatopancreas fatty acid biosynthesis was significantly affected.

Specifically, in the muscle, one of the pathways significantly negatively affected at both 24 hpi and 48 hpi is Nicotinate and nicotinamide metabolism, which concerns the metabolism of two nicotinamide coenzymes (NAD^+^ and NADP^+^) and has metabolically diverse ramification, with levels of both coenzymes being significantly reduced (Table 1; Supplementary Table 1). We have suggested that low availability of NAD^+^ was central to the metabolic changes in the muscle in response to silenced CHH gene expression, which led to reduced fluxes of glycolysis and TCA cycle and eventually to diminished ATP levels [34]. In the same vein, low availability of NADP^+^ could reduce the flux of pentose phosphate pathway (PPP), resulting in lower levels of two PPP metabolites, NADPH and ribose 5-phosphate, which have diverse metabolic functions [41, 42]. NADPH is consumed in reductive reactions including the reduction of ribonucleotides to deoxyribonucleotides, whereas ribose 5-phosphate, via PRPP (5-phosphoribosyl 1-pyrophosphate), is a precursor to nucleotide biosynthesis [43] (see Fig. 3). Thus, a negatively impacted PPP would, as a result of low NADP^+^ levels, decrease nucleotide biosynthesis, which was clearly demonstrated by significantly lower levels of nucleotides across the board (Fig. 2, 3).

**Figure 3.**
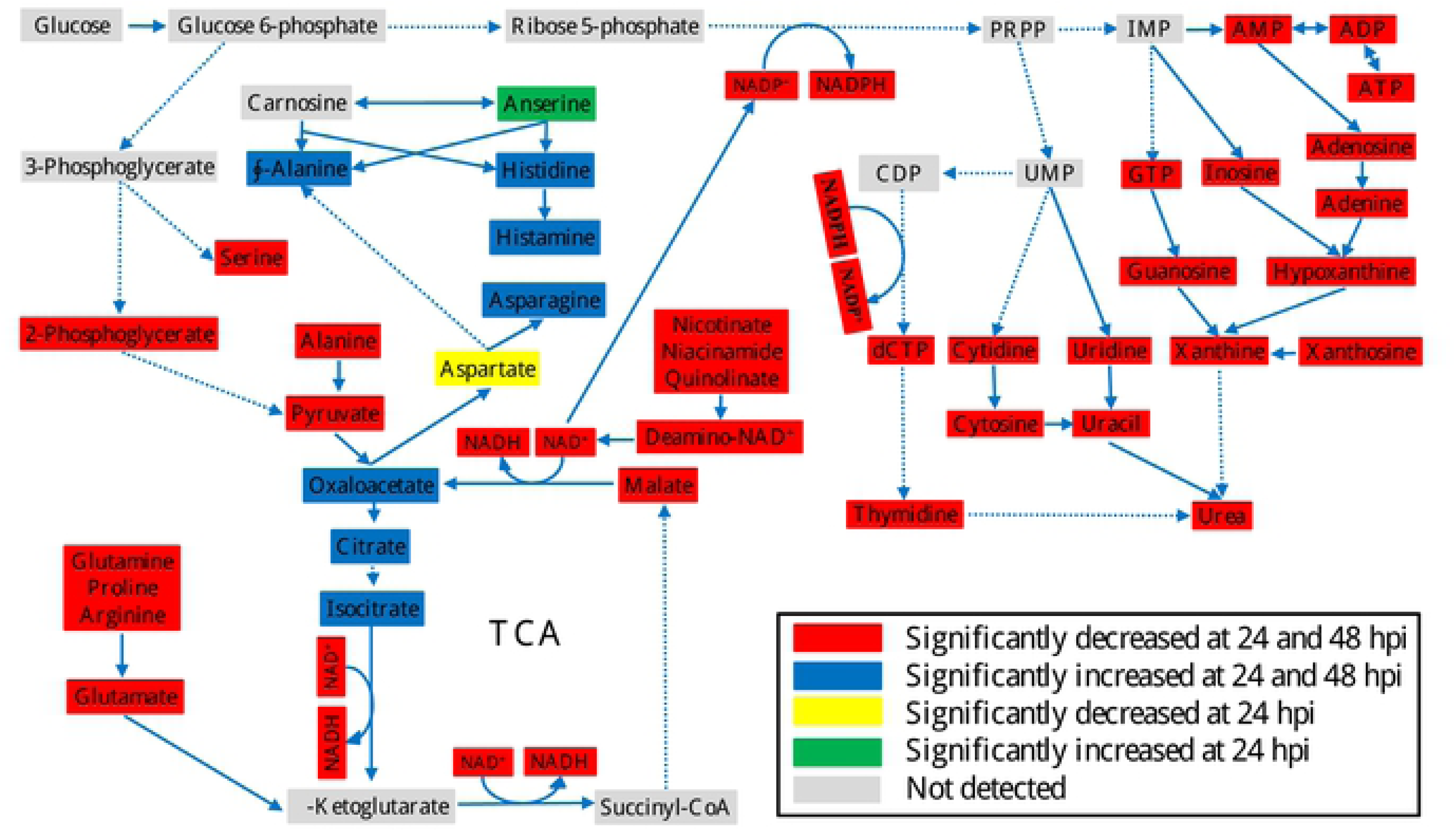
Metabolic profiles of the muscle of the crayfish *Procambarus clarkii* after CHH dsRNA treatment. Red rectangle: significantly decreased from saline-injected levels at both 24 and 48 hpi; blue rectangle: significantly increased from saline-injected levels at both 24 and 48 hpi; yellow rectangle: significantly decreased from saline-injected levels only at 24 hpi; green rectangle: significantly increased from saline-injected levels only at 24 hpi; gray rectangle: not detected. Dotted lines indicate multiple metabolic steps are involved that are not individually specified. ADP: adenosine diphosphate, AMP: adenosine monophosphate, ATP: adenosine triphosphate, CDP: cytidine diphosphate, dCTP: deoxycytidine triphosphate, Glucose 6-P: glucose 6-phosphate, GTP: guanosine triphosphate, IMP: inosine monophosphate, NAD^+^/ NADH: oxidized/reduced nicotinamide adenine dinucleotide, NADP^+^/NADPH: oxidized/reduced nicotinamide adenine dinucleotide phosphate, Ribose 5-P: ribose 5-phosphate; PRPP: phosphoribosyl pyrophosphate, UMP: uridine monophosphate.

Another pathway that was significantly affected in the muscle by CHH gene silencing at 24 hpi is Pyruvate metabolism (Table 1; Supplementary Table 1). Pyruvate sits at a site where several metabolic pathways intersect. A significantly affected pyruvate metabolism is compatible with the observations that glycolysis and TCA cycle were negatively impacted in the muscle when CHH gene expression was silenced [34]. Pyruvate could also be used to directly form alanine or converted via a TCA intermediate (α-ketoglutarate) to other non-essential amino acids such as glutamate, glutamine, arginine, proline [44], whose levels were all decreased after CHH gene expression was silenced (Figs. 1 and 3). Moreover, a glycolytic intermediate (3-phosphoglycerate) could serve as a precursor for the biosynthesis of serine [44], levels of which were significantly decreased (Figs. 1 and 3). Indeed, analyses with MetPA and MSEA confirmed that Alanine, aspartate and glutamate metabolism and several amino acid metabolite sets (Aspartate metabolism, Alanine metabolism, Glutamate metabolism, Arginine and proline metabolism, Glycine and serine metabolism) was negatively affected (Tables 1 and 3, Supplementary Tables1 and 3). On the other hand, levels of several amino acids and related metabolites in the muscle were increased after silencing CHH gene expression; these included those of asparagine, anserine, histidine, histamine, and β-alanine (Fig. 1). Aspartate and asparagine are two of the amino acids that could be synthesized from a TCA intermediate (oxaloacetate) [44], whose levels was shown to be significantly accumulated because of inhibition of the 3 NAD^+^-dependent TCA reactions that carry isocitrate through to oxaloacetate due to low NAD^+^ levels [34]. Thus, an increase in asparagine levels was likely a spillover from a TCA cycle thus inhibited (Fig. 3). In addition, increases in β-alanine, histidine, and anserine levels (Fig. 1) are worth mentioning. Carnosine (not detected) is a dipeptide, highly concentrated in the muscle and brain, consists of β-alanine and histidine, and could be converted to anserine via methylation of the histidine residue (Fig. 3) [45]. Carnosine has been shown to be able to, among others, scavenge and detoxify reactive oxygen species and reactive aldehyde derived from lipid peroxidation [45, 46]. Thus, elevated levels of β-alanine, histidine, and anserine suggest that degradation (to β-alanine and histidine) and conversion (to anserine) from carnosine were enhanced under silenced CHH gene expression and imply that CHH has protective roles for the muscle cells by increasing carnosine levels via combination of β-alanine and histidine and conversion from anserine. Finally, Glyoxylate and dicarboxylate metabolism, which was highly significantly impacted by silenced CHH gene expression (see Table 1; Supplementary Table1), describes a variety of reactions involving glyoxylate or dicarboxylates that interconnect with several aspects of cellular metabolism, including Pyruvate metabolism and TCA cycle, Glycine, serine and threonine metabolism, Purine metabolism, Nitrogen metabolism that were also impacted as revealed by MetPA and MSEA.

In the hepatopancreas, the effects of silencing CHH gene expression were comparatively milder than in the muscle. Significantly affected pathways include Pyruvate metabolism and Glycolysis or Gluconeogenesis, which are consistent with our previous suggestion that glycolysis was inhibited and gluconeogenesis stimulated [34].

MSEA analysis of the metabolites, other than confirmed affected pathways as suggested by MetPA, provided additional insights regarding the effects of silencing CHH gene expression. Thus, in the muscle, MSEA analysis identified Warburg Effect as a metabolite set being significantly negatively affected in response to CHH gene silencing at 24 hpi (see Table3 and Supplementary Table 3). An invertebrate Warburg effect, characterized by up-regulation of several metabolic pathways, including glycolysis, the pentose phosphate pathway, ribonucleotide biosynthesis, glutaminolysis and amino acid biosynthesis, was observed in the tissues of the white spot syndrome virus (WSSV)-infected shrimps [47]. Based on the tissue responses (in the muscle) to silencing of CHH gene expression, the metabolic profile of the Warburg effect induced by WSSV infection is consistent with what would be expected when CHH release is greatly enhanced. Indeed, infection of white spot syndrome virus enhanced a rapid and long-lasting release of CHH into hemolymph [48]. Thus, the combined data support the notion that the WSSV-induced Warburg effect is at least in part due to the metabolic effects of CHH released by WSSV infection. In addition, MSEA also revealed that in the muscle Ammonia recycling, which involves deamination of excess amino acids and purine nucleotides [44], was negatively impacted (see Table3 and Supplementary Table 3), which, combined with results showing significantly lower levels of amino acids and nucleotides (Figs. 1 and 2), indicated a metabolic status of low protein and nucleic acid turnovers. Moreover, analysis with MSEA indicated that in the hepatopancreas fatty acid biosynthesis was significantly affected (Table 4 and Supplementary Table 4). Due to the limitation of the methodology employed by the present study, only water soluble short-chained fatty acids (acetate, butyrate, caprate) were detected, whose levels decreased significantly (Supplementary Table 4). The results obtained under silenced CHH gene expression, though only a small sub-set of fatty acids were detected and quantified, are consistent with those from a previous study showing that CHH stimulated lipid mobilization from the storage tissues (*e.g*., the hepatopancreas), including free fatty acids release into hemolymph, to be used metabolically in the consuming tissues [49].

## 4. Conclusion

In summary, the present study, combined with methods developed in a previous study [34], established a metabolomics pipeline that includes an NMR-based detection and identification of tissue metabolites and a web-based tool suite MetaboAnalyst allowing interpretation of metabolomics data in the context of metabolic network. In conjunction with a dsRNA-based gene silencing method [34], the metabolomics pipeline characterized the metabolic effect of CHH on its target tissues as far more diverse than previously realized. In brief, CHH clearly has differential effects on the 2 target tissues, with the muscle much more heavily regulated by CHH. Whereas CHH is expected to promote carbohydrate utilization in the hepatopancreas through stimulating glycolysis and lipolysis, its stimulatory effect on nicotinate and nicotinamide metabolism plays a central role in coordinating metabolic activity in the muscle with diverse and wide-ranging consequences, including enhancing the fluxes of glycolysis, TCA cycle, and pentose phosphate pathway, leading to increased ATP supply and elevated protein and nucleic acid turnovers. Combining our data and previous ones [22, 28, 29, 31, 33, 34, 49, 50], a comprehensive scheme of CHH roles in the 2 target tissues is proposed (Fig. 4).

**Figure 4.**
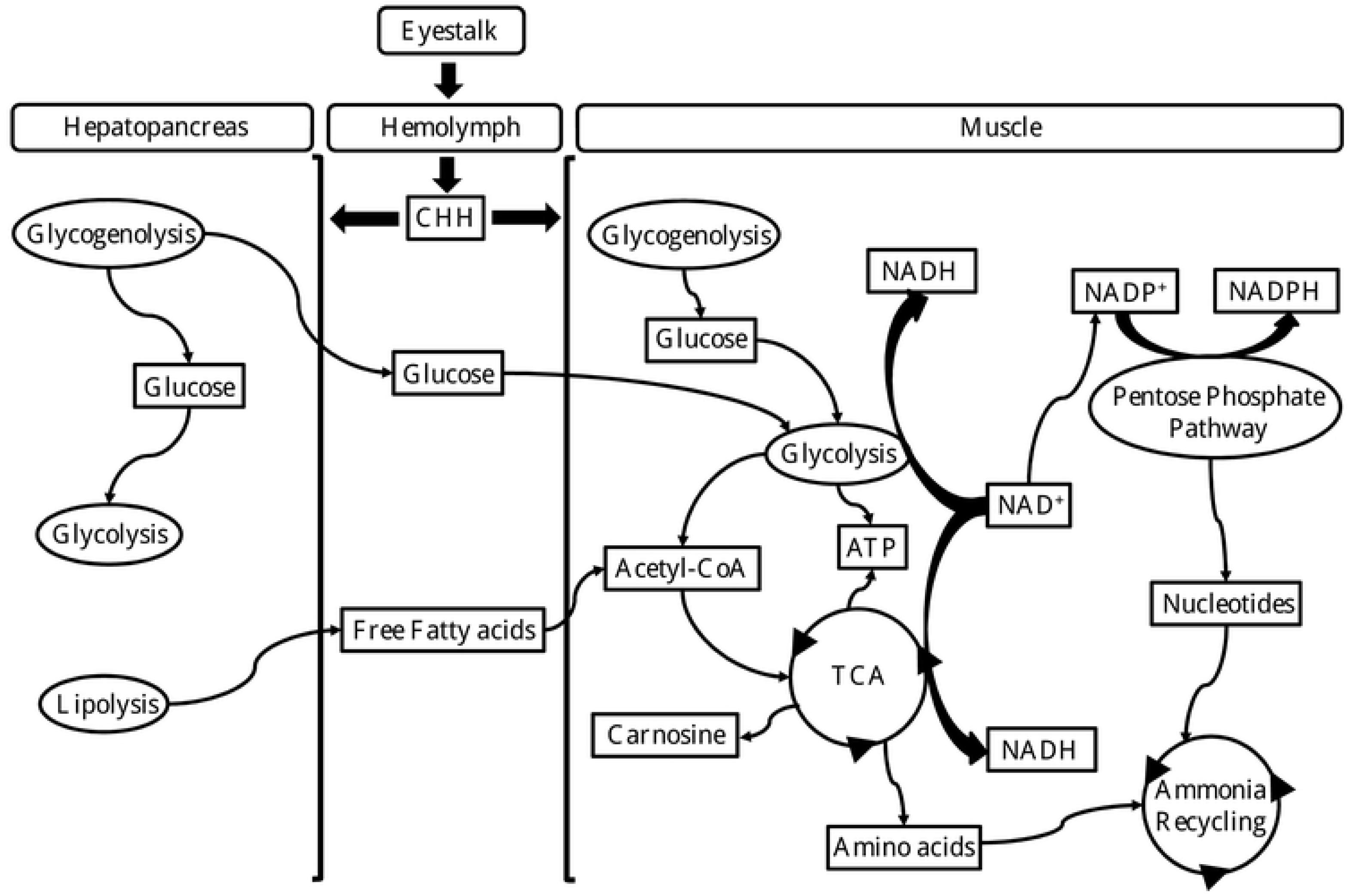
Proposed metabolic roles of CHH in the muscle and hepatopancreas. Under conditions when its release from the eyestalk ganglia into the hemolymph is stimulated (Webster, 1996; Chang *et al*., 1998; Chang *et al*., 1999; Zou *et al*., 2003; Lorenzon *et al*., 2004; Lin *et al*., 2013), CHH acts on and differentially regulates its target tissues. Hence, while CHH stimulates glycogenolysis in both target tissues, resulting in higher levels of glucose (Parvathy, 1972; Keller and Andrew, 1973; Sedlmeier, 1981, 1987; Sedlmeier and Keller, 1983) that drives glycolytic flux (Santos and Keller, 1993a,b), in the hepatopancreas, CHH additionally enhances lipolysis (Santos *et al*., 1997; the present study). Glucose and free fatty acids are released into the hemolymph and taken up by the muscle where they are further metabolized via glycolysis and TCA cycle, respectively, for ATP production. In the muscle, central to the effects of CHH is a stimulated Nicotinate and nicotinamide metabolism, which provides two nicotinamide coenzymes (NAD^+^ and NADP^+^) that drive glycolysis and TCA cycle, and the pentose phosphate pathway, respectively (Li *et al*., 2017; the present study), resulting in more ATP supply and higher protein and nucleic acid turnover. Additionally, CHH may provide protective effects to the muscle by increasing carnosine levels.

## Declaration of interest

There are no conflicts to declare.

## Acknowledgements

The authors thank Prof. Shangwu Ding, Nuclear Magnetic Resonance Core Facility at Department of Chemistry, National Sun Yat-sen University (Taiwan) for providing metabolomic analysis and related technical support. The present study was supported financially by the Ministry of Science and Technology (MOST), Taiwan through grants (104-2311-B-018-002 and 106-2311-B-018-001) to C-YL. W.L. was supported by a postdoctoral fellowship (104-2811-B-018-001) from MOST, and a China postdoctoral international exchange program and the China postdoctoral science fund (2018M632578).

## Supporting information

Supplementary Table 1 Effects of CHH dsRNA treatment on levels of metabolites involved in the KEGG metabolic pathways significantly impacted by silencing of CHH gene expression in the muscle of the crayfish *Procambarus clarkii*.

Supplementary Table 2 Effects of CHH dsRNA treatment on levels of metabolites involved in the KEGG metabolic pathways significantly impacted by silencing of CHH gene expression in the hepatopancreas of the crayfish *Procambarus clarkii*.

Supplementary Table 3 Effects of CHH dsRNA treatment on levels of metabolites involved in the SMPDB metabolite sets significantly impacted by silencing of CHH gene expression in the muscle of the crayfish *Procambarus clarkii*.

Supplementary table 4 Effects of CHH dsRNA treatment on levels of metabolites involved in the SMPDB metabolite sets significantly impacted by silencing of CHH gene expression in the hepatopancreas of the crayfish *Procambarus clarkii*.

